# Taxonomic and diet-based functional structure of carabid communities varies seasonally in cultivated fields

**DOI:** 10.1101/2023.02.14.528449

**Authors:** Caro Gaël, Marrec Ronan, Auguste Cyril, Barbottin Aude

## Abstract

Agricultural intensification has altered the provision of natural bioregulation. We assessed the effects of five different crops under non-inversion tillage on the temporal dynamics of carabid assemblages. We evaluated how the taxonomic diversity, the composition, and the diet-based functional structure of communities varied over the spring period.
Carabid assemblages were monitored over 4 years (from 2009 to 2012), in a total of 67 fields (each field followed one year) cropped with either winter oilseed rape, winter wheat, sugar beet, spring barley, or spring pea. We assigned a relative diet profile to each species accounting for more than 0.5 % of the total mean abundance-activity: granivorous, phytophagous, and zoophagous.
The seasonal changes in species richness, abundance-activity, and evenness had the same dynamic in all crops. Despite differences in species identities between crops, the zoophagous and phytophagous diet profiles showed similar temporal dynamics in all crop types, suggesting a high functional equivalence among species present in cultivated fields. Zoophagous species were progressively replaced by primarily phytophagous species in all crops, while the increase in granivorous species was only observed in oilseed rape.
Our results suggest that potential bioregulation do not significantly differ among crop types but vary along the spring season.

**Highlights:**

- We monitored carabid communities in five crop types under non-inversion tillage
- We observed similar seasonal increases in carabid abundance-activity and species richness in all crops during the spring period
- We observed similar trends in temporal changes in carnivorous and phytophagous diet profiles in all crops
- With seasonal changes, zoophagous species are progressively replaced by more phytophagous species
- Crop type is not the main factor driving temporal changes in diet-based structure of carabid assemblages

## 1. Introduction

The expansion of arable lands throughout the world and the maximization of production have threatened biodiversity in many ways (Krebs et al., 1999; Pardo et al., 2020; Tilman et al., 2001). While the expanded use of fertilizers and pesticides has increased crop yields (Tilman et al., 2002), these practices are also the cause of major losses of biodiversity and environmental issues (such as water pollution) in farmland landscapes (Geiger et al., 2010; Robinson and Sutherland, 2002; Tscharntke et al., 2005). The loss of biodiversity has affected all taxa (Donald et al., 2001), with major consequences on ecosystem functioning, and the provision of related ecosystem services such as pollination or pest regulation (Cardinale et al., 2012; Grab et al., 2018). As a response, alternative cropping systems and management practices have been proposed since the 1980s (Altieri et al., 1983; Gayer et al., 2019). Among them, diversifying and increasing the length of crop rotations have been proposed to reduce negative impacts on biodiversity, especially on ground-dwelling macroinvertebrates (O’Rourke et al., 2008).

Among important soil macroinvertebrates, carabid beetles (Coleoptera: Carabidae) are usually considered as natural enemies of pests and weeds in crop fields (Carbonne et al., 2020; Kromp, 1999; Lavelle et al., 2006; Lövei and Sunderland, 1996; Marc and Canard, 1997), and are important elements of trophic chains sustaining biodiversity (Benton et al., 2003; Thiele, 1977). An increase in either carabid abundance or species richness has been shown to potentially enhance ecosystem sustainability (Jowett et al., 2021; Kromp, 1999). For example, prey consumption by carabids was found to be positively correlated to their abundance (Menalled et al., 1999), while species richness improves community functional resilience as well as biodiversity conservation (Tilman, 1996). Carabid assemblages are also sensitive to habitat modifications (Lövei and Sunderland, 1996; Magura and Ködöböcz, 2007; Melnychuk et al., 2003; Woodcock et al., 2005), crop management (Bertrand et al., 2015; Caro et al., 2016; Holland, 2004; Marrec et al., 2015; Winqvist et al., 2011), crop rotations length (O’Rourke et al., 2008), and crop type (Marrec et al., 2015).

Because of their different sensitivities to biotic and abiotic habitat conditions, the different species in the regional pool are likely to respond differently to the environmental constraints induced by each crop type (Eyre et al., 2012; Jowett et al., 2019; Purtauf et al., 2005). For instance, cropping calendars and seasonal variations in microclimatic conditions depend on crop phenologies and alter the composition of carabid assemblages over time. These changes in carabid assemblages occurs according to specific carabid life cycles and habitat preferences. Previous studies have mainly focus on population changes and changes at a community level have been poorly studied (Marrec et al., 2015; Thomas et al., 2001). Community-level seasonal changes of species have been shown to vary between crop types (Marrec et al., 2021). Nevertheless, it is not known how such taxonomic or trait-based changes affect other aspects of functional diversity, such as pest and weed regulation potential by carabid assemblages. One way to understand the influences of these changes should be considering temporal variations of dominant diet affinities among species in carabids assemblages, using a functional trait approach. Indeed, this approach provides a direct link to mechanistic processes (e.g., type of trophic resource consumed). The likelihood of species shift in carabid assemblages tends to be correlated with the trophic level (Holt et al., 1999; Raffaelli, 2004). For example, predator carabids are considered to be characterized by a greater sensitivity to habitat changes than herbivorous species (Duffy, 2003; Holt et al., 1999; McCann, 2007; Raffaelli, 2004). However, little is known about the seasonal variation of the regulation potential associated with carabid species simultaneously present in different crop types usually found in agricultural landscapes. Thus, a study combining both taxonomic and functional approaches may provide food for thought to design crop rotations and landscape-level management practices aiming at supporting key ecosystem services such as pest control (Tscharntke et al., 2012).

Here, we aim at exploring how the taxonomic and diet-related functional diversity of carabid beetle communities varied during spring in response to crop type. We used data from a monitoring design conducted over four years in a total of 67 fields from 2009 to 2012 and in five crop types: winter wheat, winter oilseed rape, sugar beet, spring barley, and spring pea. These five crops have been selected to maximize the differences in terms of cropping calendars (i.e., influencing plant phenology and practices) and vegetation density and structure (i.e., influencing microclimate conditions). Both parameters are also likely to impact food resource availability. More specifically, we address the following hypotheses: (i) similar seasonal dynamics of taxonomic and functional diversities of carabid communities are expected in all crop types as all crops change in their vegetation structure in the time period and (ii) diet-based composition of carabid communities is expected to change differently during the season depending on crop type.

## 2. Materials and methods

### 2.1 Sampling sites

The study was conducted every year from 2009 to 2012 during four consecutive months each year in sites located in Seine-et-Marne, France (80 km southeast of Paris). The area is dominated by cereal crop production managed conventionally (in average for the four years within 500m around the survey fields: 45 % of winter wheat, 15 % of sugar beet, 11 % of winter oilseed rape, 8% of spring barley, 7 % of legumes, and 2 % of maize or sunflower). The carabid assemblages were monitored in a total of 67 fields—each one being sampled only one single year (mean field size ± SD: 11 ± 5.6 ha). Fields cropped with the five dominant crops in the study area were monitored: winter oilseed rape (*n* = 9 fields), winter wheat (*n* = 27 fields), sugar beet (*n* = 12 fields), spring barley (*n* = 11 fields), and spring pea (*n* = 8 fields) (see Table 1 for details). Because of its high prevalence in the landscape, winter wheat was the most frequent crop sampled. As tillage practices have a significant impact on ground-dwelling arthropods (Hatten et al., 2007; Kosewska et al., 2014; Shearin et al., 2014), we standardized this practice and only selected sampled fields conducted under non-inversion tillage for at least five consecutive years before the sampling. Study fields were conventionally managed with mineral fertilizers and chemical pesticides. Agricultural practices in each field were recorded based on farmers’ interviews (Table 1). Studied crops highly differ in their sowing dates and pesticide pressure expressed as Treatment Frequencies Index (TFI), which includes all the types of treatments (Butault et al., 2011; Pingault et al., 2009; Sattler et al., 2007) (Table 1).

**Table 1:** Main characteristics of the farming practices for each crop monitored over the five years

| | Number of fields sampled | Fields' surface area (ha)(mean ± sd) | Sowing date (earliest – latest) | Nitrogen rate (kgN/ha) (mean ± sd) | TFI* total (mean ± sd) | TFI* insecticide (mean ± sd) |
| --- | --- | --- | --- | --- | --- | --- |
| Winter wheat | 27 | 12.2 ± 6.2 | 16 Oct. – 22 Oct. | 186 ± 30 | 4.87 ± 1.53 | 0.95 ± 0.81 |
| Sugar beet | 12 | 12.0 ± 6.5 | 18 March – 22 March | 122 ± 18 | 6.00 ± 1.12 | 0.51 ± 0.65 |
| Oilseed rape | 9 | 10.7 ± 4.2 | 26 August – 2 Sept. | 173 ± 30 | 5.82 ± 1.80 | 3.27 ± 1.55 |
| Spring barley | 11 | 10.4 ± 4.8 | 19 Feb. – 06 March | 126 ± 16 | 2.90 ± 0.61 | 0.31 ± 0.54 |
| Spring pea | 8 | 9.3 ± 4.7 | 27 Feb. – 04 March | 0 | 3.85 ± 0.92 | 1.72 ± 0.70 |

\* Pesticide use intensity was estimated using the Treatment Frequency Index (TFI) (Butault et al., 2011; Pingault et al., 2009). The TFI was estimated as the ratio between the amount of pesticide applied in a field per hectare, and the recommended rate per hectare. Two types of TFI were calculated, namely TFI insecticide and the total TFI, estimated as the sum of the TFI of all pesticides used.

### 2.2 Carabid sampling

Carabids were sampled using three pitfall traps per field and then we averaged the three abundance-activities. When using pitfall traps, some limitations may arise when comparing mean abundance-activity (MAA) between crops due to differences in capture probability (Knapp et al., 2020; Lang, 2000; Thomas et al., 2006). Nevertheless, this method remains widely used in most studies on ground-dwelling arthropods. In addition, we are primarily interested in the temporal variation in MAA in each crop type individually, which limits the potential impact of this bias.

The first trap was placed at approximately 30 m from a field edge and the other traps were placed at least 60 m from each other and from the edge. Pitfall traps were filled with a saturated salt (NaCl) solution and a few drops of detergent and protected from litter and rainfall by a lid (15 cm × 15 cm). The traps were installed once every month for four months, from March to June (except in 2009, from April to June), and kept open for 7 consecutive days. We focused on the spring period because it is a period of high activity for carabids and a critical period for crop pest management. Following trap collection, carabids were stored in the lab in a 70 % ethanol solution and identified at the species level following Jeannel (1942, 1941). Species names follow *Fauna Europaea* (Jong et al., 2014).

### 2.3 Statistical analyses

All statistical analyses were performed in R 3.5.0 (R Core Team, 2018).

Carabid communities in crops are usually numerically dominated by a few species, which may drive agro-ecosystem functioning (Holland and Luff, 2000; Luff, 2002; Smith and Knapp, 2003). To avoid an effect of rare species in analyses of compositional and functional diversity, only species representing at least 0.5 % of the total MAA were included in subsequent analyses (Mazzia et al., 2015; Pelosi et al., 2014) (see ESM 1 for the list of selected species).

To avoid intra-field variance and the bias due to pseudo-replication, we calculated the total MAA for each sampling date and field as the average number of captured carabid individuals from the three pitfall traps. We also calculated the species richness and Pielou’s evenness index (*J’*) (Krebs, 1989) per field, considering only the subset of dominant species. *J’* measures the distribution of individuals within each species assemblage, irrespective of the species richness, and ranges from 0 (dominance of one single species) to 1 (equal proportion of all species in the assemblage).

#### Effect of crop type and sampling date on taxonomic indices

For all subsequent models, we used generalized linear models (GLM) with a Poisson distribution and log-link function when considering the species richness or abundance-activity as the response variable, and GLM with a quasi-binomial distribution and log-link function for the evenness index. To test for absolute differences between crops we ran independent GLMs for each taxonomic index (*n* = 3) using the crop type (*n* = 5 levels) as the explanatory factor. To investigate seasonal trends in community composition in each crop type, we ran independent GLMs for each taxonomic index using the date of sampling (expressed as annual Julian days) in interaction with the crop type as explanatory variables.

#### Seasonal variations of species identity composing carabid assemblages

The seasonal variation in the species composition of carabid assemblages of the most abundant species was characterized using multivariate analyses. We performed a Principal Component Analysis (PCA) using the MAA of each species listed in Table 2 in every community. We used a VARIMAX procedure to maximize the correlation between PCA axes and the MAA of each species. We selected PCA axes with eigenvalue > 1 and percentage of total explained inertia > 10 %. Then, we recorded the coordinates of each species along all selected PCA axes. To evaluate if community composition changed significantly over season, we used linear models (LMs) to test the effect of the date of sampling (expressed as annual Julian days) on the PCA coordinates of each community (i.e., species assemblage at a given sampling location and date). Analyses were performed with the R package ADE4 (Dray et al., 2007).

**Table 2:** Diet profile of the carabid species representing at least 0.5% of the total abundance. For each diet modality (Zoophagous, Phytophagous and Granivorous), we calculated the proportion represented by each modality in the species’ diet profile according to the literature. For each carabid species, we provided the number of references where we found information about their diet.

| Carabid species | Zoophagous | Phytophagous | Granivorous | Number of references (Total = 82) |
| --- | --- | --- | --- | --- |
| *Amara similata* | 0,12 | 0,31 | 0,57 | 16 |
| *Anchomenus dorsalis* | 0,57 | 0,33 | 0,1 | 7 |
| *Brachinus explodens* | 1 | 0 | 0 | 2 |
| *Harpalus affinis* | 0,24 | 0,31 | 0,44 | 26 |
| *Loricera pilicornis* | 0,85 | 0,15 | 0 | 6 |
| *Nebria brevicollis* | 0,92 | 0 | 0,08 | 14 |
| *Notiophilus biguttatus* | 1 | 0 | 0 | 13 |
| *Poecilus cupreus* | 0,58 | 0,25 | 0,17 | 27 |
| *Pseudoophonus rufipes* | 0,33 | 0,33 | 0,35 | 12 |
| *Pterostichus melanarius* | 0,4 | 0,51 | 0,1 | 24 |
| *Pterostichus niger* | 0,75 | 0 | 0,25 | 8 |

#### Crop type and sampling date effect on diet-based functional indices

To inform the diet profile of all selected abundant species (Table 2) we compiled information from 82 references (See ESM 3 for details). Diet profiles were defined according to the distribution of the food preferences of each species identified in the literature in three main diets: granivorous, phytophagous, and zoophagous. Given the heterogeneity of data, we homogenized information using the fuzzy coding method (Chevene et al., 1994) following the procedure described in Pelosi et al. (2014). Information was coded by an affinity score ranging from 0 to 3 for each of the three main diets, and affinities were summed up to build the trait profile (e.g., the distribution of affinities within diets). We standardized the affinities for each diet in order that the sum of the affinities for a given species equals 1 (see Table 2). Finally, we calculated the community-weighted means (CWM) for each diet separately. CWM reflects the mean affinity value to a diet of the community weighted by the relative MAA of each species (Garnier et al., 2004; Lavorel et al., 2008; Violle et al., 2007) (hereafter denominated as CWM_granivorous_, CWM_phytophagous_, and CWM_zoophagous_).

To test pair-wise correlations between the CWMs in each crop type, we used GLM with a binomial distribution and log-link function (as CWM was bounded between 0 and 1). We used models with one CWM as response variable and the two other CWM as quantitative variables in interaction with the crop type.

To investigate seasonal trends for the three CWM related to diet profiles in each crop type, we ran independent models for each CWM indices (*n* = 3) with the sampling date (expressed as annual Julian days) in interaction with the crop type as explanatory variables. We used a GLM with a binomial distribution and log-link function for the three CWM related to diet profiles.

## 3. Results

During the four years of sampling, we collected 81,968 individuals, corresponding to 56 carabid species. The quantities of beetles trapped each year have been highly variable due to the contrasting weather conditions. Eleven carabid species had a relative abundance-activity of at least 0.5 %, representing 80,861 individuals (Table 2, ESM 1 & 2). When considering all years and crops together, *Pterostichus melanarius*, *Poecilus cupreus*, and *Anchomenus dorsalis* were the three most abundant sampled species, accounting for 43.3, 30.7, and 7.1 % of the total captures, respectively.

### 3.1 Crop type and sampling date effect on taxonomic indices

Over the entire sampling period, winter crops (*i.e*., oilseed rape and wheat) had a significantly higher species richness (respectively, 6.1 ± 2.1 and 5.2 ± 1.8 species among our subset of species) compared with spring crops (*i.e*., sugar beet, pea and spring barley, respectively 4.2 ± 1.5, 4.4 ± 1.7 and 4.6 ± 1.4 species) (*p* < 0.05; Figure 1). The MAA was significantly higher in oilseed rape than in the other crops (*p* < 0.05; Figure 1). Evenness index was higher in oilseed rape and wheat than in sugar beet (respectively, 0.6 ± 0.16, 0.56 ± 0.20 and 0.47 ± 0.24, *p* < 0.05; Figure 1) while spring barley and pea had intermediate values (respectively 0.52 ± 0.20 and 0.51 ± 0.20, *p* < 0.05; Figure 1).

**Figure 1:**
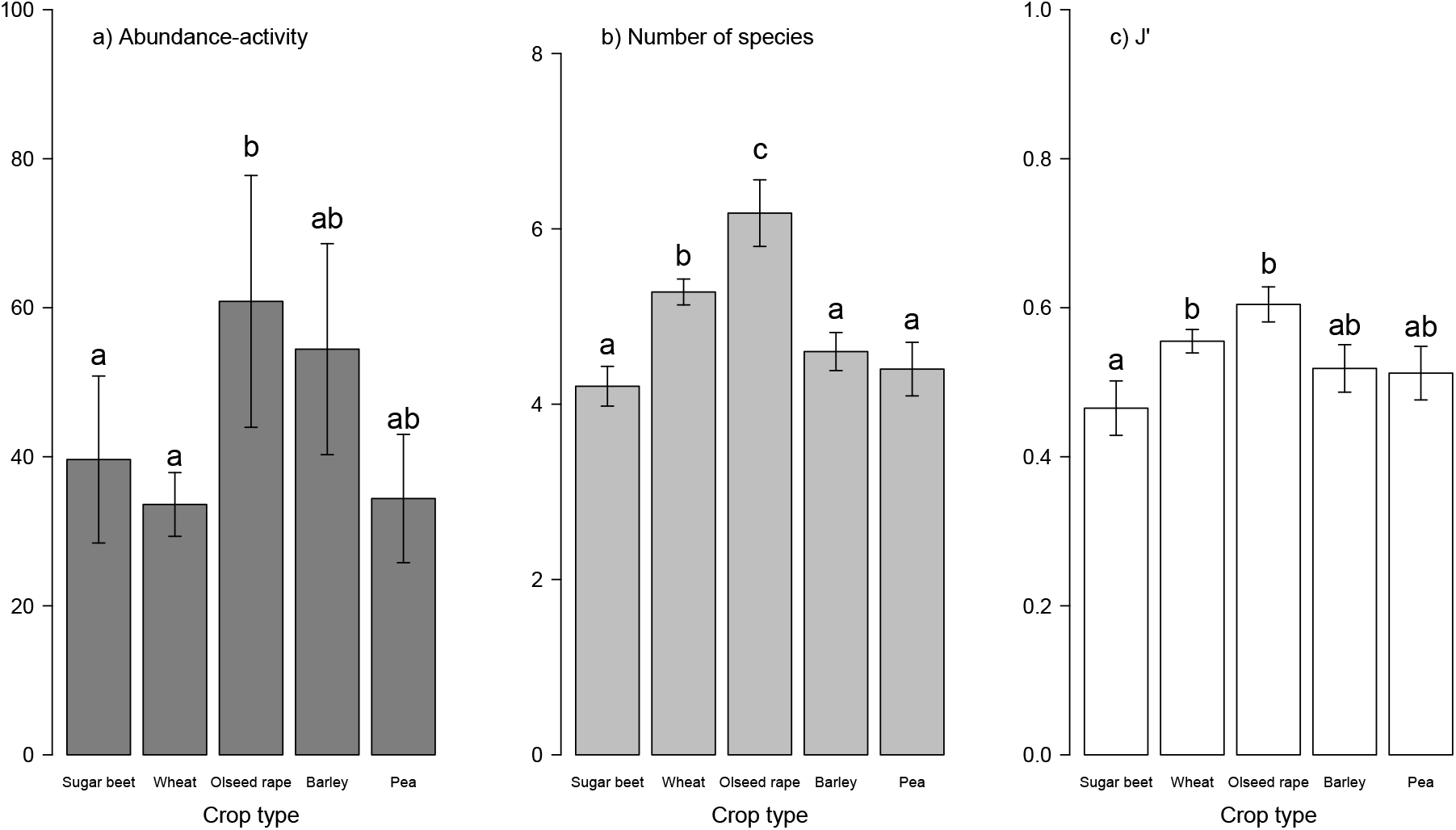
a) Abundance-activities, b) number of species and c) evenness index of dominant species of carabids (mean ± standard error) for each crop. Each different letter represents a significant difference in statistical analyses (GLM with a Poisson distribution and log-link function for the species richness and the abundance-activity, GLM with a quasi-binomial distribution and log-link function for the evenness index).

In all crops, MAA increased significantly during spring along the sampling season expect in pea crops (Figure 2). Considering the ratio between June and March, in descending order, MAA heightened by 14.90 in sugar beet, by 12.35 in oilseed rape, by 7.47 in spring barley and by 2.50 in wheat. Concerning species richness, it increased significantly in all crops expect in spring barley (Figure 2). Considering the ratio between June and March, in descending order, species richness heightened by 1.52 in pea, by 1.40 in oilseed rape, by 1.23 in sugar beet and by 1.1 in wheat. Surprisingly, the evenness did not change during the same period, except in sugar beet where it halved, considering the ratio between June and March (Figure 2).

**Figure 2:**
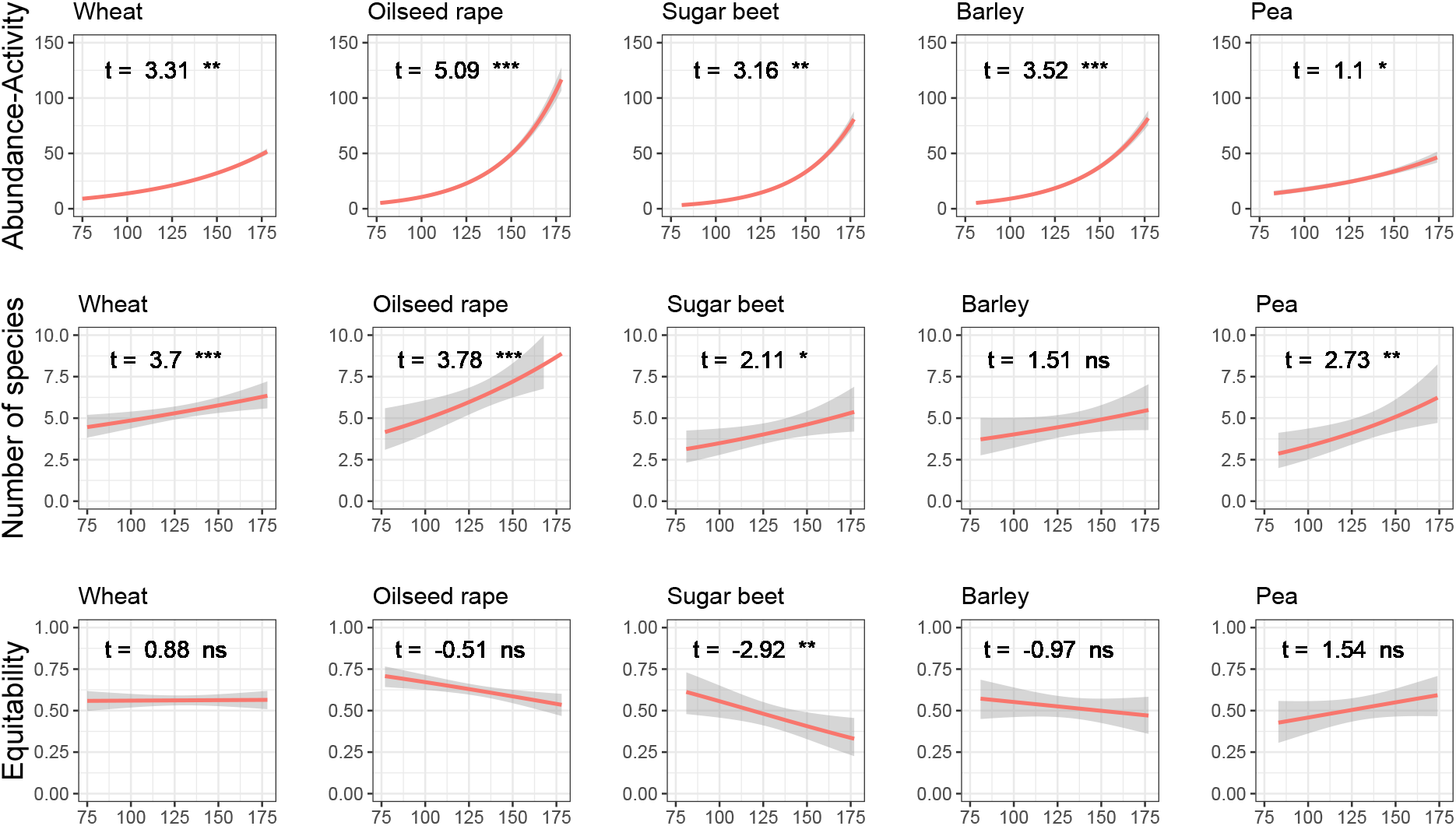
Change in abundance-activity (first line), number of species (second line) and evenness index (third line) along time (Julian days) for each crop (considering the dominant species of carabids). The red line represents linear regression using GLM with poisson family for abundance-activity and number of species and GLM with quasi-binomial family for equitability. The t-value and the significance (illustrated by stars or *ns* for *non-significant*) are generated by the same models (for each indice, one model with crops and julian days in interaction as explanatory variables). The shaded area represents the standard error.

### 3.2 Seasonal variations of species identity composing carabid assemblages

The Principal Component Analysis showed that several species discriminated assemblages along three leading dimensions (total variance explained: 54 %; Figure 3). The first PCA axis (28 %) was positively associated to MAA of *Amara similata*, *Brachinus explodens*, and *Harpalus affinis*. The second PCA axis (16 %) was negatively associated to MAA of *Notiophilus biguttatus* and *Pterostichus niger* and positively to the one *P. melanarius* and *Pseudoophonus rufipes*. Finally, the third PCA axis (10%) was positively associated to MAA of *Nebria brevicollis*. The MAA of both *P. cupreus* and *A. dorsalis* characterised in the same way the three PCA axes, suggesting that these two species were equivalently present in all crops during the entire monitoring.

**Figure 3:**
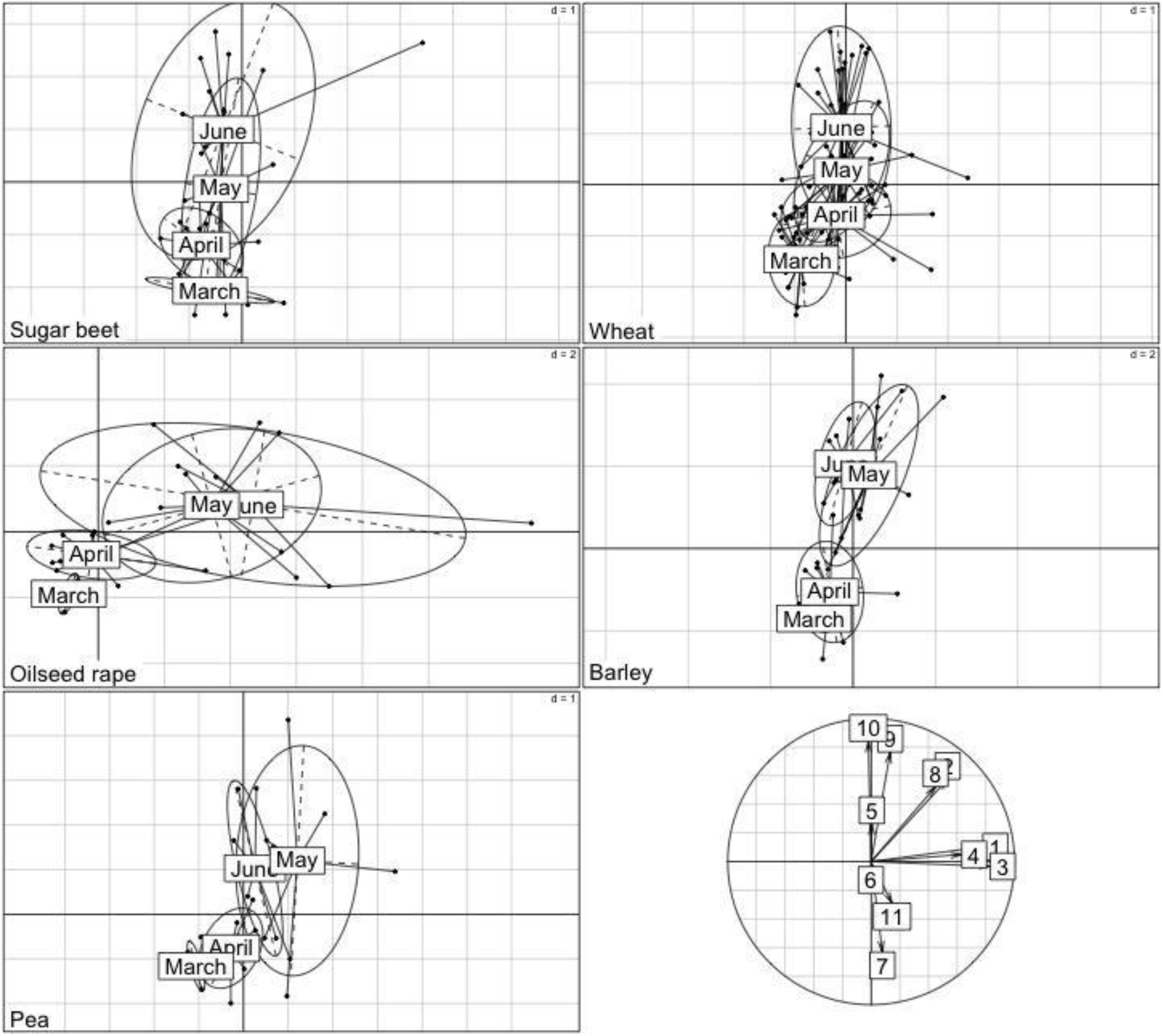
Ordering of the dominant species of carabid assemblages according to their composition in species in the plane defined by the PCA axis 1 and 2 for the five crops sugar beet, wheat, oilseed rape, barley and pea. We ran one PCA but decided to separate graphically the 5 types of crop in order to be clearer. Each ellipse represents a month (March, April, May or June). In the correlation circle, each number represents one species: 1: *A. similata*, 2: *A. dorsalis*, 3: *B. explodens*, 4: *H. affinis*, 5: *L. pilicornis*, 6: *N. brevicollis*, 7: *N. biguttatus*, 8: *P. cupreus*, 9: *P. rufipes*, 10: *P. melanarius*, 11: *P. niger*.

Once species assemblages were allocated to their PCA coordinates, we detected a strong temporal shift in species composing carabid assemblages during the sampling period. Except for sugar beet, the weighted coordinates of species assemblages on the three PCA axes were significantly influenced by the Julian date in all crop types (*p* < 0.01). Concerning sugar beet, we only detected a significant effect of time along the second PCA axis. In March, all five crops were mainly characterized by *N. biguttatus* and *P. niger*. Later in the season, the carabid assemblages were gradually characterised by *P. rufipes* and *P. melanarius* in wheat, barley and sugar beet and by *A. similata*, *B. explodens* and *H. affinis* in oilseed rape and, in a least extent, pea.

### 3.2 Crop type and sampling date effect on diet-based functional indices

The CWM_zoophagous_ and CWM_phytophagous_ were significantly and negatively correlated in every crop type (*p* < 0.05). We found that the functional assemblages in the early season were mainly characterized by species primarily zoophagous as illustrated by the higher values of CWM_zoophagous_ at this period (Figure 4). CWM_zoophagous_ decreased significantly during spring along the sampling period in all crops except in pea crops (Figure 4). Considering the ratio between June and March, in descending order, the CWM_zoophagous_ decreased by 0.88 in pea, by 0.85 in wheat, by 0.74 in spring barley, by 0.71 in sugar beet and by 0.57 in oilseed rape. The reverse dynamic was observed for the CWM_phytophagous_, which increased during spring in all crops (Figure 4). Considering the ratio between June and March, in descending order, the CWM_phytophagous_ heightened by 1.97 in sugar beet, by 1.77 ion spring barley, by 1.41 in wheat and by 1.37 in both oilseed rape and pea. These results suggest that, as the spring season progresses, carabid assemblages are gradually dominated by species with a dominant phytophagous diet profile (Figure 4). On the other hand, the CWM_granivorous_ was very low, indicating that predominantly granivorous species are never dominant in studied carabid assemblages (Figure 3). Nevertheless, the relative proportion of such species increased significantly during spring in oilseed rape (increasing by 2.09) whereas it decreased significantly in wheat crops (decreasing by 0.97) (Figure 4).

**Figure 4:**
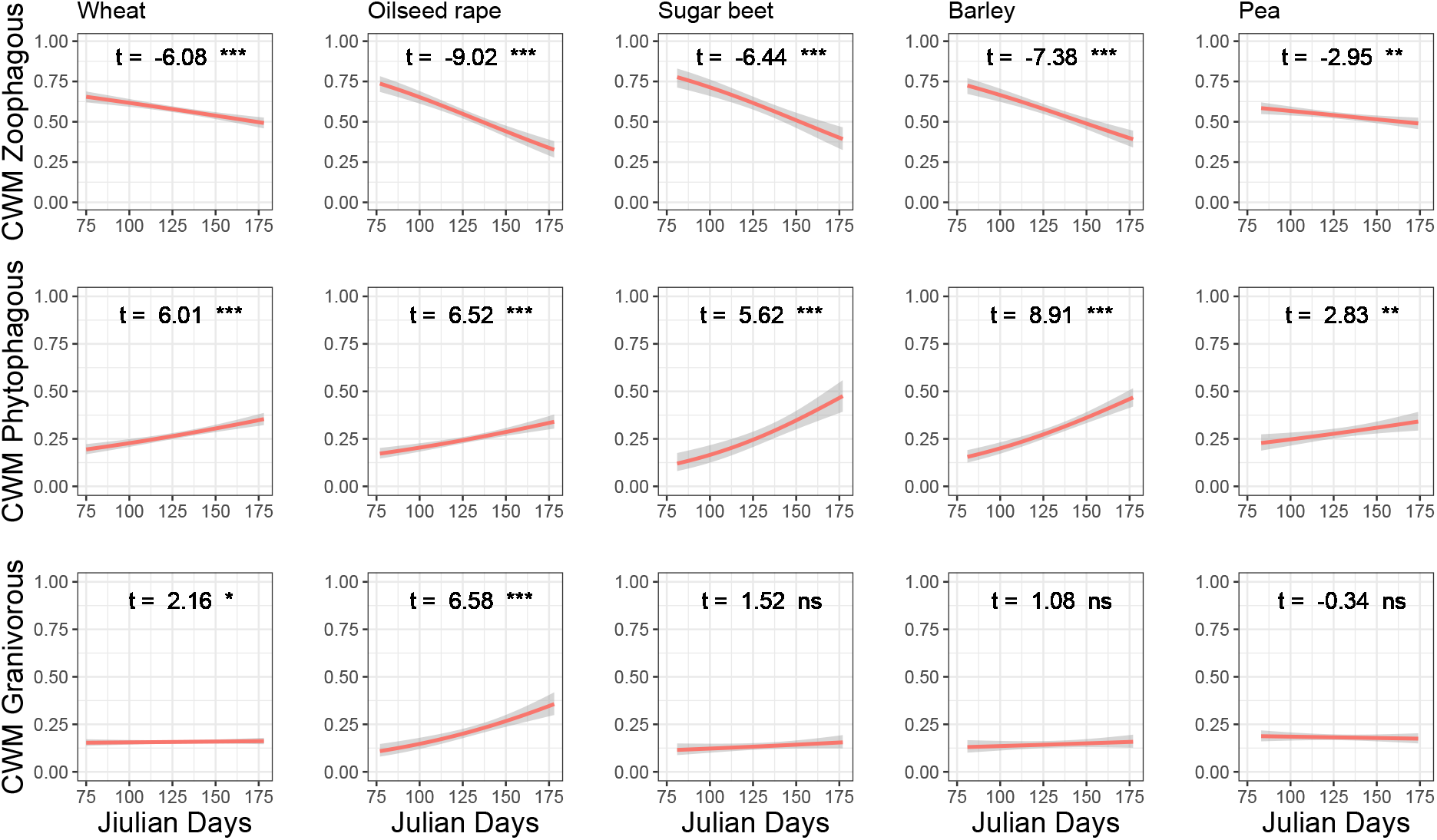
Change in the three Community Weighted Means (CWM) related to diet over time (Julian days) based on dominant species of carabids. The red line represents linear regression using GLM with quasibinomial family. The shaded area represents the standard error. The t-value and the significance (illustrated by stars or *ns* for *non-significant*) were generated by the same models (GLM with quasibinomial family).

## 4. Discussion

In this study, we explored how the taxonomic and diet-based functional diversities of carabid beetle communities inhabiting various crop types varied during spring. We have shown that the taxonomic structure and species composition of carabid assemblages varies across spring season and between crop types. Indeed, we observed that the taxonomic changes in community assemblages (in terms of abundance-activity, species richness, or evenness) had similar temporal trends in the five studied crops (Figure 2), congruently with our first hypothesis. This congruence is particularly interesting knowing that species identity and relative abundance-activity varied importantly among crops (Figure 3). In addition, despite these differences in species composition, the diet-based functional assemblages and their temporal trends showed similar trends in all crop types (Figure 4). This result indicates that the potential for biological control provided by dominant carabid species assemblages had similar temporal trend across the season, independently of crop type, but the amount of variation was crop dependant (Begg et al., 2017).

### Crop type as a main species filter

Carabid abundance-activities and species richness increased significantly during spring in all crops. This is consistent with the findings of Holland and Reynolds (2003) and Knapp et al (2019) and many other studies on ground-dwelling invertebrates. A possible reason is that individuals are more active as the season progresses because of the increase in ambient temperature, thereby influencing the abundance-activity. As other macroinvertebrates, all carabid species have specific phenologies which affect the timing of their emergence as adults during the spring season (Matalin, 2007). Consequently, more and more species are susceptible to appear in carabid assemblages during spring. In addition, some species may colonize crop fields during spring and therefore increase species pool within field during this period (Holland et al., 2004; Marrec et al., 2015; Thomas et al., 2002). On the other hand, evenness remained stable in all crop types during spring. This index considers both the relative abundance-activity of each species in the assemblage and the total number of species. Although the species richness increased over the season, the stability of the evenness illustrates that the relative abundance-activity of each species decreased within the carabid assemblages, but communities remain as unevenly dominated all along the season. The processes behind this stable pattern remain unclear and should be the subject of future research. For instance, knowledge on intra-guild competition and use of available ecological niches within agricultural fields is notably lacking on ground-dwelling generalist predators, including carabid beetles, but also spiders or rove beetles.

As illustrated by the multivariate analysis (Figure 3), species composition in local assemblages differed between crop types and from month to month, with some species associated with specific crops at specific times. Crop types differed in terms of farming practices (Table 1), timing of cultivation (Table 1), and vegetation structure, which can act as environmental habitat filters affecting the composition of carabid assemblages (Booij and Noorlander, 1992; Hance et al., 1990; Holland, 2004). Oilseed rape was one of the most treated crops, with insecticides applied mainly in spring (April and May, mean TFI_insecticide_ = 3.27 ± 1.55; Table 1). But it is also the crop with the most developed vegetation in spring compared to spring crops, which could explain its greater attraction for ground beetle communities (Hanson et al., 2017; Seidl et al., 2020). Spring barley was the crop receiving the least pesticide treatments (mean TFI = 2.90 ± 0.61). As for spring pea, it is sown from the last days of February to the first decade of March (Table 1). Sugar beet was the last crop sown in the studied fields, during the last decade of March, and received high levels of herbicides in spring (TFI = 6.00 ± 1.12; Table 1). Winter wheat was the main crop cultivated in the region, and it is sown in October and holds a high variability of pesticide levels (TFI = 4.87 ± 1.53; Table 1). In summary, each carabid species may respond differently to combinations of practices (Weibull et al., 2003); differences in fertilization are likely to generate different communities of weeds, which are consumed by carabids (Pyšek and Lepš, 1991); different types of crops are likely to attract different pests, which may be consumed by carabids (Bebber et al., 2014); and different sowing dates generate an environment that may be attractive to carabid species to varying degrees (Marrec et al., 2017, 2015). In their review, Holland and Luff (2000) discussed in depth the impact on carabids of treatments such as pesticides, sowing date, and fertilization. They concluded that many factors governed carabid species assemblages and that, despite the large number of studies, it was still difficult to understand which factors governed their distribution. Consequently, the combination of all these influences should generate specific carabid species assemblages for each crop type, which is what we found.

### Diet-based functional dynamic is similar between crop types

One novelty of our study was to consider the diversity in diet affinities of each carabid species and not considering one trophic type for each species (Table 2). Thanks to this approach, we found that the relative dominance of zoophagous and phytophagous diet profiles presented similar temporal trends during the spring season among all local assemblages, independently of the type of crop. Carabid assemblages were progressively characterized by primarily phytophagous species (such as *P. rufipes* and *P. melanarius)* to the detriment of predominantly zoophagous species (such as *N. brevicollis, P. niger* and *N. biguttatus*). Carabids are well known for being consumers of crop pests (Kromp, 1999; Lövei and Sunderland, 1996) which have specific phenologies. Depending on the period of the year, availability of trophic resources changes. Carabid emergence begins from February and lasts until summer (Purvis and Fadl, 1996). The emergence or the presence of animals is often synchronous with the one of their resources (de Souza Mendonça Jr, 2001). In February, the main resources available might be crop pests that have not yet emerged or seeds. Eggs remaining in the soil or larvae are available in this period and are easy to catch (Lövei and Sunderland, 1996; Thiele, 1977). This prevalence of animal resources should explain the dominance of zoophagy in our results (Figure 4). After preys become adults, they might start moving and become more difficult to catch compared to seeds, or they could disperse outside the crops in response to crop management (pesticide use, mechanical weeding…). Weeds germinate in fields mainly during spring and therefore become more available for phytophagous species during this season (Hartzler et al., 1999; Milberg and Andersson, 1997; Roberts and Pottier, 1980). Regarding the granivorous diet expression, we observed a significant seasonal increase only in oilseed rape, suggesting the presence of more species mainly granivorous (such as *A. similata*, *B. explodens* and *H. affinis*) than in any other crops in late spring. This trend is in accordance with previous work (González et al., 2020). At the end of June, oilseed rape is one of the crops with the most mature seeds and thus that have the highest probability of falling to the ground. In addition, Deroulers et al. (2019) showed that the lipid content is one of the main drivers of seed consumption by carabid beetles. As oilseed rape seeds are one of the richest seed in lipid (Friedt and Snowdon, 2010), this characteristic may explain the preferred use of oilseed rape by granivorous species.

Although we did not quantify the resource availability for carabids, numerous studies observing prey availability (animals and weeds) supported our hypothesis (de Souza Mendonça Jr, 2001; Hartzler et al., 1999; Lövei and Sunderland, 1996; Milberg and Andersson, 1997; Roberts and Pottier, 1980; Thiele, 1977). Nevertheless, antagonist biological control effects are never to be excluded, because of variation in species efficacy and interspecific interactions (Letourneau et al., 2009). Negative interactions, such as intraguild predation (Finke and Denno, 2004; Julian et al., 2019) or interference (Charalabidis et al., 2019), can disrupt the efficiency of prey depletion (De Heij and Willenborg, 2020).

## 5. Conclusion

We have shown that abundance-activity and species richness increased during spring in five types of crops managed in non-inversion tillage. Changes also occur in the species composition and community differ between crops during spring. However, these changes led to the same temporal trend in diet-based functional structure between the 5 crop types studied. This functional shift was probably a response to a change in resources available for carabids. Annual crops in agroecosystems show a high variability in their phenology and management, depending on their type. These differences between crop types likely affect their abiotic characteristics and the availability of biotic resources during spring. Based on our observations, crop types and presumably associated management practices did not seem impacting either taxonomic (in terms of abundance-activity and evenness of species assemblages) or diet-based functional temporal trends of carabid assemblages but only the identity of dominant species in carabid assemblages. To maintain bioregulation in agricultural landscape, crop diversification should be combined to other solutions benefiting to all the biodiversity providing biocontrol service (directly and indirectly) and not only the carabids. Thus, we emphasised the complementarity of a mixed taxonomic and diet-based functional approach in agroecological studies and the necessity to consider the temporal scale when evaluating the potential for biological control. In addition, our study highlighted the necessity to consider the identity of carabid species composing communities and the diversity in their diet affinities and not only synthetic community indices in order to improve natural pest control by maximizing the presence of species functionally benefiting to crop protection.

## Acknowledgments

The ANR - Agence Nationale de la Recherche (The French National Research Agency) provided partial funding to this research, under the “SYSTERRA programme - Ecosystems and Sustainable Development", as part of the “ANR-08-STRA-007, FARMBIRD - Coviability models of FARMing and BIRD biodiversity” project. We are grateful to Clémence Bouty, Bertrand Couillens, Solène Orrière and Catherine Cresciucci from the UMR SADAPT research unit who helped with field sampling and species identification. We also thank Colin Van Reeth and Jodie Thénard from UMR 1121 (University of Lorraine) for valuable discussions and for reading the manuscript.

## Appendix

ESM 3: List of the 82 references used to inform the diet affinities of 11 carabid species selected in this study

---

**ESM 1:**
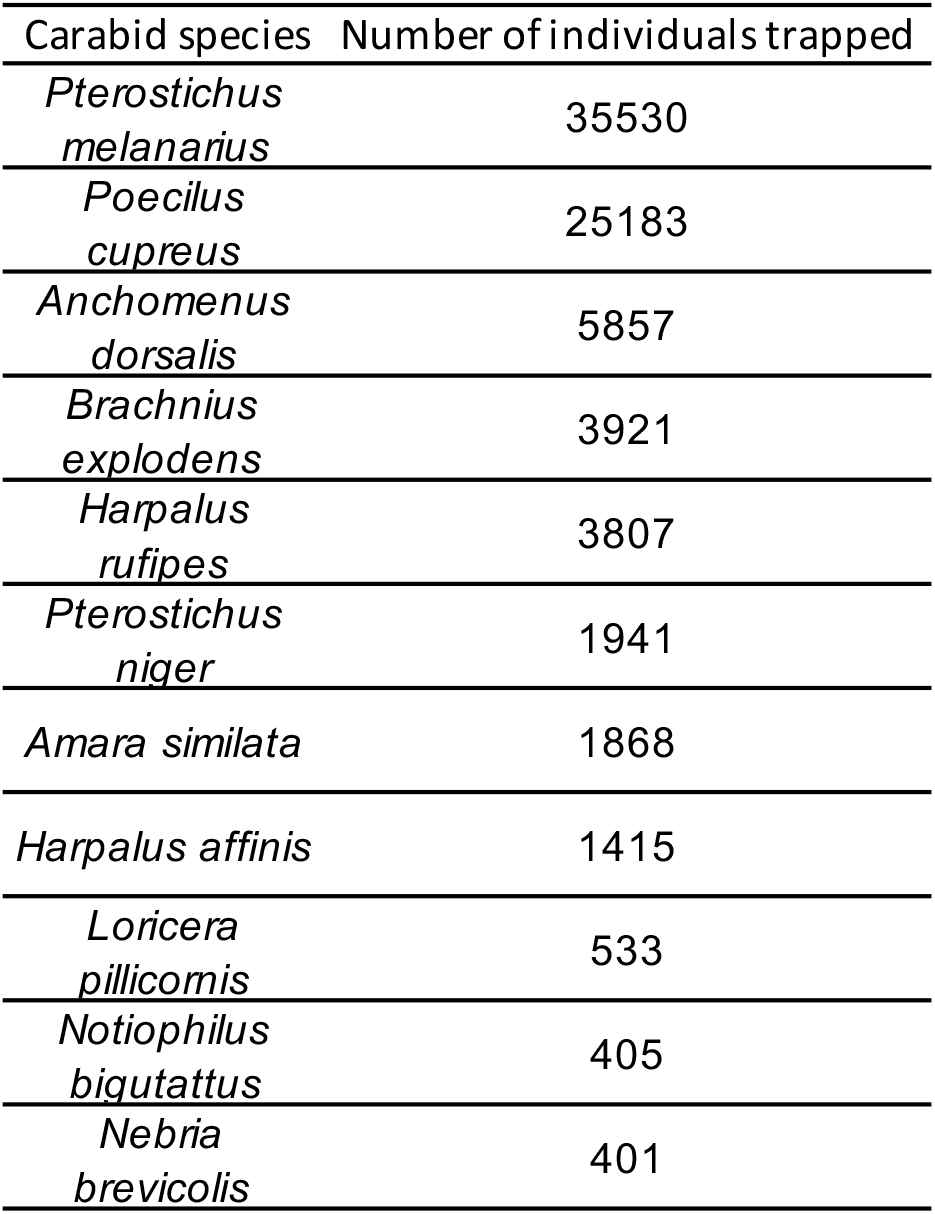
Abundance-activity of the 11 dominant carabid species representing each at least 0.5 % of the total abundance-activity.

**ESM 2:**
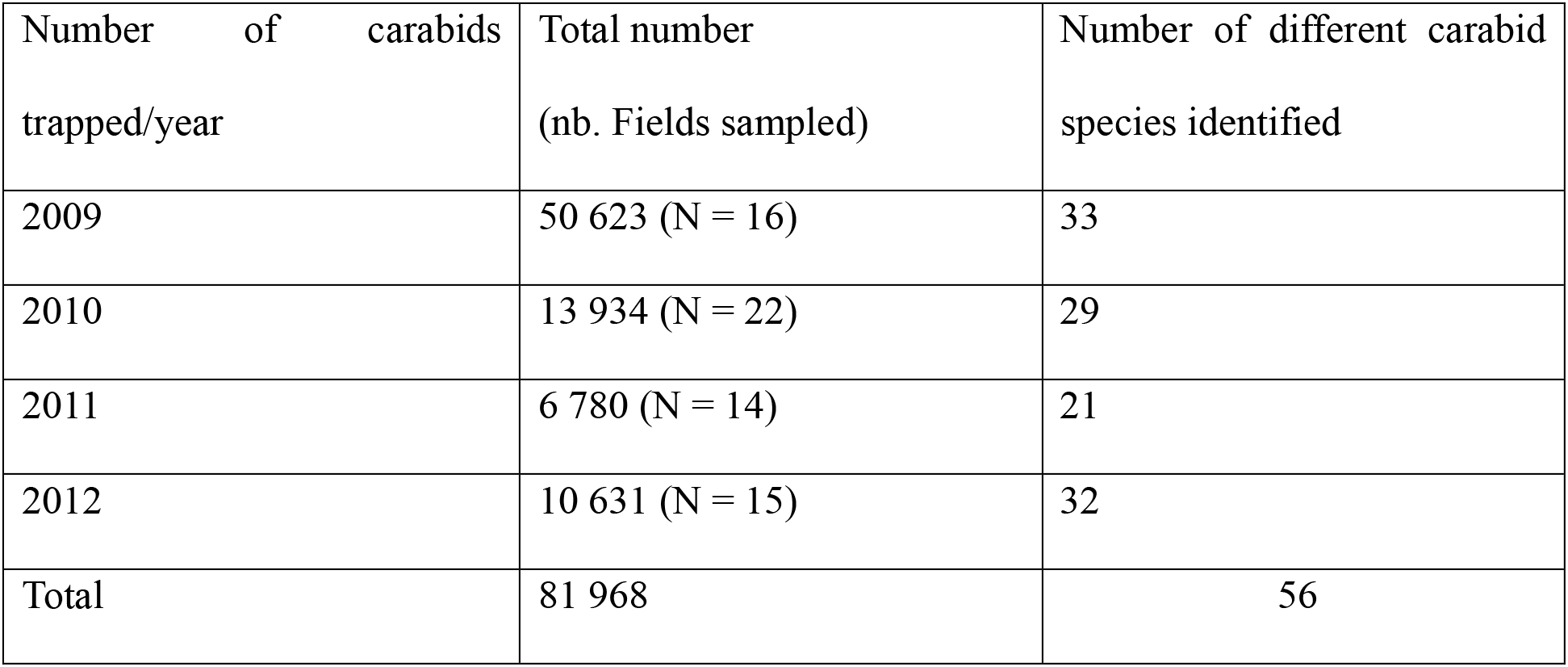
Total number of carabids trapped per year, in bracket, total number of fields sampled each year and number of different carabid species identified each year.

